# Endosperm evolution by duplicated and neofunctionalized Type I MADS-box transcription factors

**DOI:** 10.1101/2021.06.08.447573

**Authors:** Yichun Qiu, Claudia Köhler

**Affiliations:** Department of Plant Biology, Uppsala BioCenter, Swedish University of Agricultural Sciences and Linnean Centre for Plant Biology, Uppsala, 75007, Sweden; Max Planck Institute of Molecular Plant Physiology, Potsdam Science Park, Am Mühlenberg 1, 14476 Potsdam-Golm, Germany

## Abstract

MADS-box transcription factors (TFs) are present in nearly all major eukaryotic groups. They are divided into Type I and Type II that differ in domain structure, functional roles, and rates of evolution. In flowering plants, major evolutionary innovations like flowers, ovules and fruits have been closely connected to Type II MADS-box TFs. The role of Type I MADS-box TFs in angiosperm evolution remains to be identified. Here, we show that the formation of angiosperm-specific Type I MADS-box clades of Mγ and Mγ-interacting Mα genes (Mα*) can be tracked back to the ancestor of all angiosperms. Angiosperm-specific Mγ and Mα* genes were preferentially expressed in the endosperm, consistent with their proposed function as heterodimers in the angiosperm-specific embryo-nourishing endosperm tissue. We propose that duplication and diversification of Type I MADS-genes underpins the evolution of the endosperm, a developmental innovation closely connected to the origin and success of angiosperms.

## Introduction

MADS-box transcription factors (TFs) are an evolutionary ancient class of TFs and major developmental regulators present in nearly all major eukaryotic groups (Alvarez-Buylla et al. 2000). They have largely amplified during land plant evolution and play important roles in regulating organ patterning and timing of reproductive developmental programs (Nam et al. 2003; Gramzow and Theißen 2013). The loosely conserved DNA binding MADS-domain is located at the N-terminus of MADS-box proteins, while based on the C-terminal sequences two types of MADS-box TFs are distinguished, Type I and Type II (Alvarez-Buylla et al. 2000; Schwarz-Sommer et al. 1990). The duplication and divergence of Type II MADS-box genes, or MIKC-type, has been linked to the evolution of floral organs in angiosperms, including flowers, ovules and fruits (Nam et al. 2003; Becker and Theissen 2003; Ruelens et al. 2013; Ruelens et al. 2017). Compared to Type II, Type I MADS-box genes are underrepresented in gymnosperms and have experienced more frequent lineage-specific duplications in angiosperms, followed by fast pseudogenization and gene loss (Nam et al. 2004; Gramzow and Theißen 2013). Nevertheless, the role of Type I MADS-box TFs in angiosperm evolution remains to be identified. Emerging studies suggest a role for Type I MADS-box genes in the regulation of female gametophyte and endosperm development in *Arabidopsis* and grasses (Bemer et al. 2008; Colombo et al. 2008; Steffen et al. 2008; Shirzadi et al. 2011; Roszak and Köhler 2011; Hehenberger et al. 2012; Chen et al. 2016; Batista et al. 2019; Zhang et al. 2020; Paul et al. 2020). In this study, we establish a link between the evolution of Type I MADS-box genes and the origin of the endosperm in flowering plants. We hypothesize that through gene duplication and neofunctionalization, novel subfamilies of Type I MADS-box TFs acquired endosperm-specific function in the shared common ancestor of all extant angiosperms after its divergence from gymnosperms. This process likely underpinned the evolution of the endosperm in angiosperms.

## Results and Discussion

### Duplication of Mβ and Mγ MADS-box TF genes is concerted with the evolution of angiosperms

We identified Type I MADS-box genes in 40 species, representing all major lineages of angiosperms and other land plants as outgroups (Supplementary Table S1). The phylogeny of Type I MADS-box genes in all angiosperms revealed three major clades (Fig. 1; Supplementary Figure S1), corresponding to the previously defined groups Mα, Mβ and Mγ (Parenicová et al. 2003; Arora et al. 2007; Gramzow and Theißen 2013). Specifically, we found Mγ type genes in all angiosperms we assayed (Supplementary Table S2), including *Amborella trichopoda*, the species earliest-diverging from other angiosperms, suggesting the presence of an ancestral Mγ MADS-box gene in the most recent common ancestor of all angiosperms. Mβ genes in angiosperms are sister to the angiosperm Mγ clade, while the most closely related homologs in three major lineages of gymnosperms, *Picea abies*, *Ginkgo biloba*, and *Gnetum luofuense* (previously identified as *Gnetum montanum* in the genome project) (Wan et al. 2018; Hou et al. 2020), form a clade that is the outgroup of the angiosperm Mγ / Mβ clade, followed successively by Mβ-like genes in the fern *Salvinia cucullata*, the clubmoss *Selaginella moellendorffii* and the mosses *Physomitrella patens* and *Sphagnum fallax*. Supporting previous findings (Gramzow et al. 2014), ancestral seed plants probably possessed only Mβ-like genes, in form of pre-duplicated Mβ/γ genes (Fig. 1). After the divergence from the ancestral gymnosperms, a gene duplication event in the common ancestor of all angiosperms gave rise to the Mγ clade, thus most likely there was at least one ancestral angiosperm Mβ gene and one ancestral angiosperm Mγ gene inherited in all the descendant lineages of angiosperms (Fig.1).

**Fig. 1.**
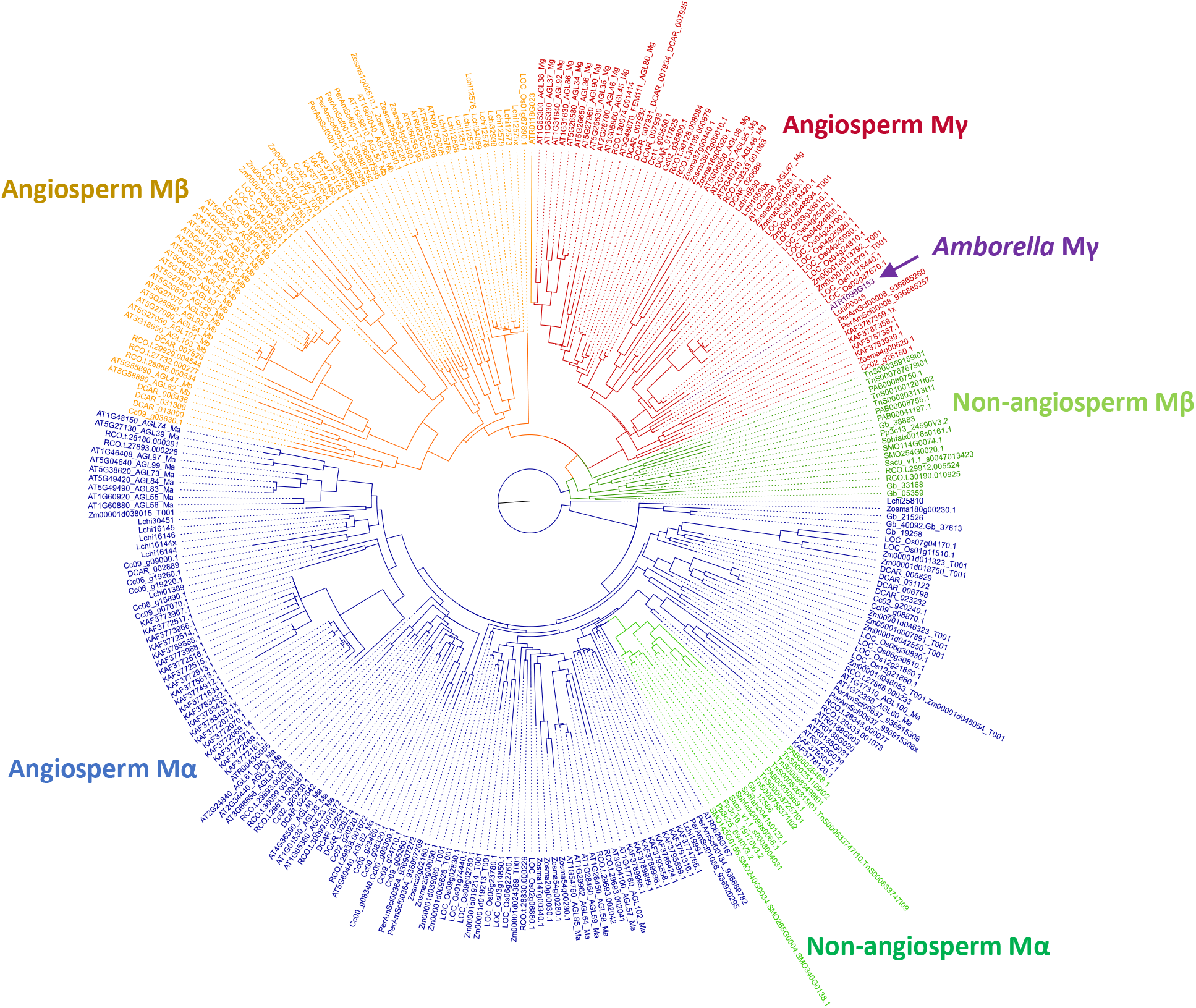
Phylogeny of Type I MADS-box TFs in selected land plants. Gene identifiers as in Supplementary Table S1-2.

### Expression of Mγ MADS-box TF genes in the endosperm is ubiquitous across the phylogeny of angiosperms

We investigated the expression patterns of the duplicated Type I MADS-box genes to pinpoint their regulatory roles in certain tissue types. Based on transcriptome data across different organs and developmental stages in *Arabidopsis thaliana* (Klepikova et al. 2016), Mγ genes were preferentially expressed in seeds and siliques, but rarely in vegetative tissues (Fig. 2a). Using available microarray data from dissected seed tissues (Belmonte et al. 2013), we inferred that several Mγ genes were mainly expressed in the early developing endosperm, but less or absent in the other compartments of seeds, such as seed coat or embryo (Fig. 2b). These data suggest that Mγ MADS-box TFs have endosperm-specific functions in *A. thaliana*. Consistent with this notion, the Mγ MADS-box TF PHERES1 is a master regulator of a gene regulatory network controlling endosperm development (Batista et al. 2019).

**Fig. 2.**
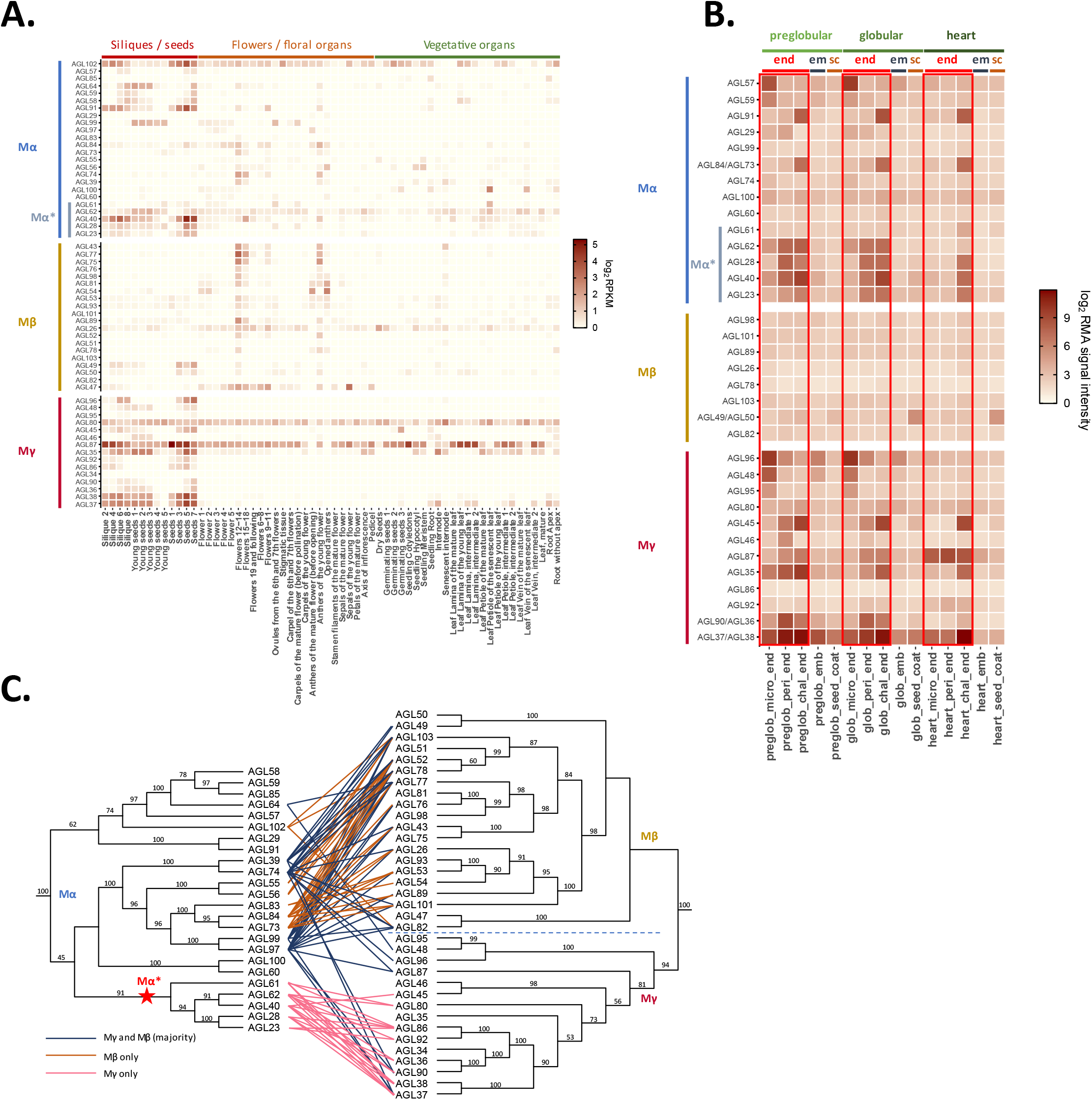
Expression of Type I MADS-box genes and interaction of Type I MADS-box proteins in *Arabidopsis thaliana*. A. Expression of *Arabidopsis* Type I MADS-box genes across different organ types and developmental stages (data from Klepikova et al. 2016). B. Expression of *Arabidopsis* Type I MADS-box genes across seed tissue types and developmental stages (data from Belmonte et al. 2013). end: endosperm; em: embryo; sc: seed coat; micro: micropylar; peri: peripheral; chal: chalazal; preglob: preglobular; glob: globular. C. Interaction between *Arabidopsis* Mα TFs and Mγ or Mβ TFs (based on yeast-two-hybrid data from De Folter et al. 2005 and Bemer et al. 2010). Phylogeny of Type I MADS-box genes is shown as ML trees with bootstrap values supporting the branches.

We also investigated the endosperm transcriptomes of maize, coconut, castor bean, soybean, and tomato and found at least one of the Mγ genes to be expressed in the endosperm of each species, consistent with their proposed roles in endosperm development (Fig.3). The Mγ genes of maize and soybean had either none or minimal expression in the embryo, supporting an endosperm-specific function (Supplementary Figure S2). Mγ gene expression was also detected in whole-seed transcriptomes of rice, avocado and monkeyflower (Fig.3). Since the orthologous Mγ genes were primarily expressed in the endosperm in other species, we infer that the observed Mγ expression in whole-seed transcriptomes likely reflects transcription predominantly in the endosperm. Thus, Mγ genes are ubiquitously expressed in the endosperm of various species representing major lineages of angiosperms, including eudicots, monocots and magnoliids, indicating that endosperm expression of Mγ genes is a conserved feature of angiosperms.

**Fig. 3.**
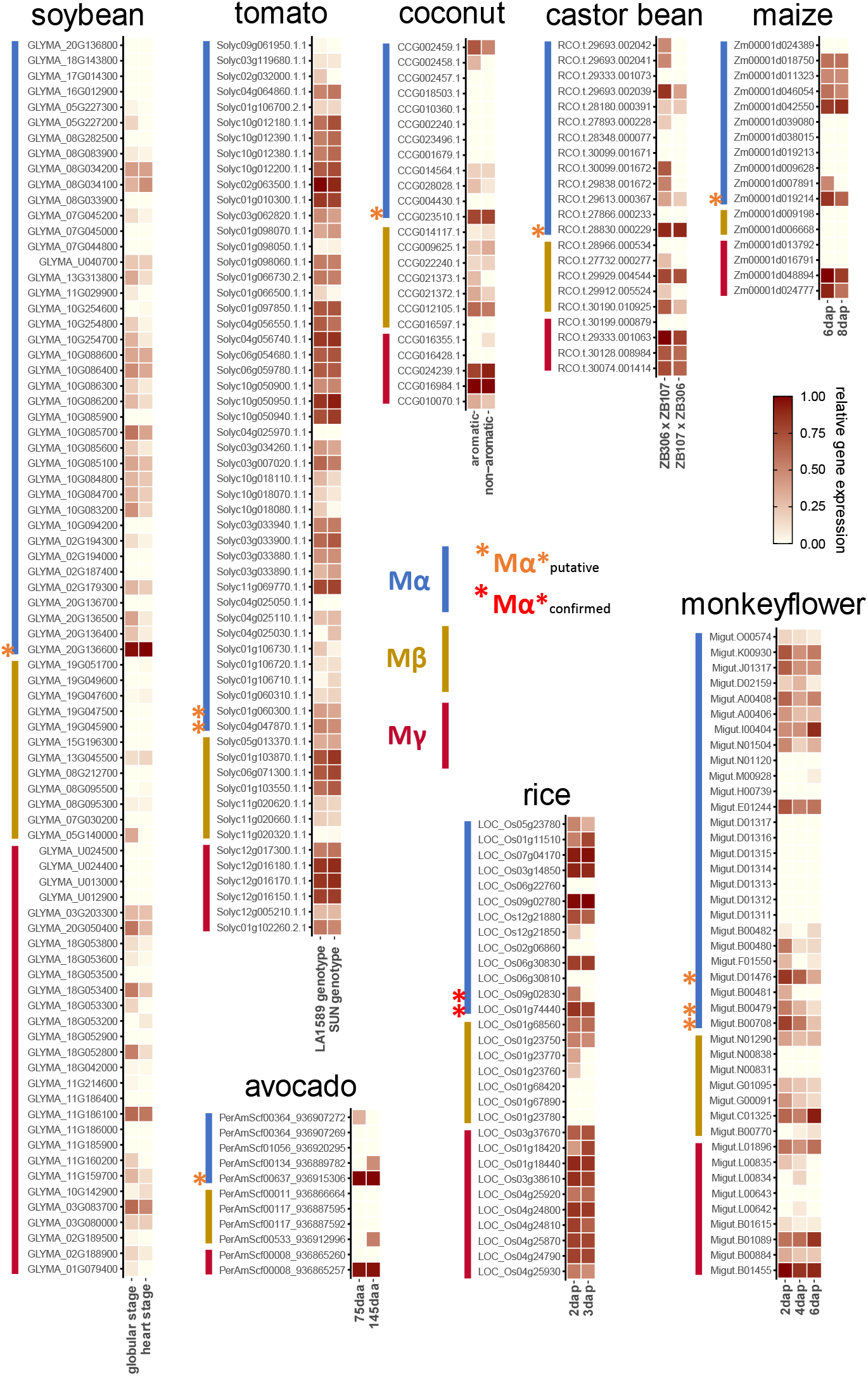
Expression of Type I MADS-box genes in flowering plants. Upper panels: endosperm transcriptomes of tomato, soybean, coconut, castor bean and maize. Lower panels: whole-seed transcriptomes of avocado, rice and monkeyflower. In each panel the expression values were normalized into a 0-1 spectrum, with the max value set as 1. For soybean, maize, avocado and rice, gene expression levels at two developmental stages; for monkeyflower, three stages are shown; dap: days after pollination; daa: days after anthesis. For tomato and coconut, gene expression levels in two genotypes (LA1589 / *SUN*) or varieties (aromatic / non-aromatic) are shown. For castor bean, gene expression levels in two reciprocal crosses between lines ZB306 and ZB107 are shown.

In contrast, Mβ genes in *A. thaliana* were barely expressed in the endosperm or other seed tissues, only one of them had low expression in the seed coat (Fig. 2b). Similarly, in maize transcriptomes, Mβ expression was not detected in the endosperm (Fig. 3). While Mβ expression was detectable at variable levels in the endosperm transcriptomes of coconut, soybean, castor bean and tomato (Fig. 3), the expression level of Mβ genes was substantially lower compared to the corresponding Mγ expression. Based on whole-seed transcriptomes, Mβ genes in avocado were nearly not expressed, Mβ genes in rice were expressed at low level, while some Mβ genes in monkeyflower were active at later stages of seed development compared to Mγ genes. The sporadic occurrence of Mβ gene expression in the endosperm or other seed tissues across the phylogeny of angiosperms suggests that the function of Mβ is dispensable in the context of endosperm regulation. In support of this notion, Type I MADS-box genes with known functional roles in the endosperm are either Mγ or Mα type genes (Bemer et al. 2008; Colombo et al. 2008; Steffen et al. 2008; Shirzadi et al. 2011; Roszak and Köhler 2011; Hehenberger et al. 2012; Chen et al. 2016; Batista et al. 2019; Zhang et al. 2020; Paul et al. 2020). Absence of Mβ genes was previously reported for the orchids *Apostasia shenzhenica*, *Phalaenopsis equestris* and *Dendrobium catenatum*, and the loss of Mβ genes was proposed connected to the deficiency of endosperm in orchids (Zhang et al. 2017). Nevertheless, some orchid species undergo double fertilization and form a rudimentary endosperm (Pace 1907; Sood and Mohana Rao 1988), suggesting that loss of Mβ is not directly related to the loss of endosperm formation in orchids. In agreement with this view, transcripts of Mα and Mγ are present in developing seeds of *A*. *shenzhenica* and *P*. *equestris* (Zhang et al. 2017), likely derived from the arrested endosperm. In *A. thaliana*, expression of some Mβ genes could be detected in the female gametophyte (Bemer et al. 2010), raising the hypothesis that their functional role is restricted to maternal tissues, rather than the endosperm. We tested this hypothesis by investigating the transcriptomes of species with perispermic seeds, in which the maternally-derived perisperm rather than the endosperm provides nutrients to the embryo. Consistent with the proposed functional role of Mβ genes in maternal tissues, we detected Mβ transcripts in the transcriptome assembly from perisperm of *Coffea arabica*. Likewise, in *Nymphaea thermarum* perispermic seeds, transcript levels of Mβ genes were much higher compared to the barely detectable Mγ gene transcripts (Supplementary Fig. S3), consistent with the perisperm accounting for the majority of the seed volume in *Nymphaea* (Povilus et al. 2015). We also investigated transcriptomes of gymnosperm reproductive tissues to infer the functional role of pre-duplicated Mβ/γ orthologs (Supplementary Fig. S3). Mβ/γ orthologous genes were expressed in female cones of *Picea abies* and ovules of *Gnetum luofuense* and expression of some Mβ/γ orthologous genes could also be detected in developing seeds of *Gnetum luofuense*, suggesting these genes perform important roles in the maternal reproductive tissue and possibly regulate the maternal nourishing behaviour supporting the development of seeds. In gymnosperms, the large female gametophyte nourishes the embryo after fertilization; while in angiosperms this role has been adopted by the endosperm which develops alongside the embryo after fertilization (Baroux et al. 2002). Based on our data we propose that the function of pre-duplicated Mβ/γ genes was to control nutrient provisioning in the female gametophyte, a function that is maintained by angiosperm Mβ genes acting in the female gametophyte and perisperm, while Mγ genes neofunctionalized and adopted an endosperm-specific function, likely enabling endosperm development.

### Duplication of Mα genes and specialization of interaction with Mγ and Mβ

MADS-box TFs usually form homo-or heterodimers (Kaufmann et al. 2005). In *A. thaliana*, an atlas of MADS-box interactions based on yeast two-hybrid data revealed distinct interaction patterns between Type II and Type I TFs (de Folter et al. 2005). Type II TFs can homodimerize, and usually heterodimerize only with other Type II TFs. In contrast, Type I TFs unlikely form homodimers, nor do they heterodimerize within the Mα, Mβ and Mγ subgroups. Instead, Mα TFs interact with members of the Mβ and Mγ subgroups, while Mβ TFs and Mγ TFs barely interact, consistent with their intrinsic relatedness compared to Mα TFs. Notably, we found that the Mα TFs (AGL62, 40, 28, 23, 61) that mainly interact with Mγ TFs clustered in a single clade (Fig. 2c). Another cluster contained Mα TFs that interact specifically with Mβ TFs and Mα TFs that have the potential to interact with both, Mβ and Mγ. Genes encoding for the obligate Mγ-interacting Mα TFs (Mα* hereafter) were specifically expressed in reproductive tissues and co-expressed with *PHE1/2* and other genes encoding for Mγ TFs in the endosperm (Fig. 2). In contrast, genes encoding for Mβ-interacting Mα TFs, as well as Mβ genes were not expressed in the endosperm. Those Mα TFs that were able to interact with both, Mβ and Mγ TFs, did not co-express with Mγ TFs in the endosperm, making it unlikely that they are able to form functional heterodimers with Mγ TFs.

We next investigated if there are Mα TFs specialized to be Mα* in other angiosperms. A *bona fide* Mα* TF is expected to have central cell / endosperm-enriched or endosperm-specific expression and interacts with Mγ TFs. Based on these predictions, *Arabidopsis* AGL62, 40, 28, 23, 61 classify as Mα* TFs. In rice, the Mα type TFs MADS78 and 79 interact with the Mγ type TFs MADS87 and MADS89 and the interaction between the two Mα TFs and Mγ TFs is required for endosperm development (Paul et al. 2020). We found that the two rice Mα genes are closely related with each other in the same subclade (Fig.1; Fig. S1). Knockout of both, *MADS78* and *79* genes, results in endosperm failure and seed lethality (Paul et al. 2020), revealing that other Mα TFs that putatively interact with Mβ TFs cannot complement the Mγ-interacting function in the endosperm.

To test whether the functional divergence of Mα genes can be detected in other angiosperm species, we analyzed the expression of Mα genes in the transcriptomes of endosperm or seeds where Mγ expression could be detected. We also found Mα genes to be highly expressed specifically in the endosperm or seeds in those species, suggesting that the regulatory divergence between the Mα* genes and other Mα genes took place across the angiosperm phylogeny (Fig. 3). We hypothesize that in response to the duplication of Mβ and Mγ genes, the duplicated Mα genes specialized in protein-protein interactions and subsequently the novel interacting pairs, Mα* and Mγ, together occupied the endosperm regulatory niche.

While the phylogeny of Mα group Type I MADS-box TFs in land plants was difficult to resolve, there is only a single cluster of Mα genes in non-flowering plants (Fig. 1). Thus, the Mα-like genes in non-flowering plants have not undergone the diversification observed in angiosperms, so they likely represent the ancestral interacting partners of the pre-duplicated Mβ-like genes (Fig. 1). In contrast, several rounds of duplications gave rise to angiosperm-specific Mα TF clades that could diverge to Mα* genes (Fig. 1), in concert with the duplication of Mβ and Mγ clades.

We observed that many angiosperm species have at least two clusters of divergent Mα genes, including the early-diverging groups *Amborella* and *Nymphaea*. Furthermore, the Mα gene phylogeny of all major angiosperm groups is largely, although imperfectly, reflected by a two-clade pattern, despite the uncertainty at the basal nodes with quite short branches (Fig.1; Fig. S1). A parsimonious model to describe the evolution of Mα type genes in angiosperms is that ancestral angiosperms most likely already possessed two, if not multiple types of Mα genes that arose from angiosperm-specific duplication. These could then diverge by forming heterodimeric complexes with either Mβ or Mγ interacting partners. Supportive to this model, we observed that in all the eudicot species we surveyed, there are Mα genes closely related to the AGL62 clade of *Arabidopsis* and expressed in the endosperm or seed transcriptomes; likewise, the expressed Mα genes in maize and coconut are in the same clade as MADS78/79 of rice. These putative Mα* genes likely derived from the same Mα* origin. Alternatively, it is also possible that several events of Mα* specialization took place convergently in angiosperms. In summary, we conclude that duplication of Mα genes and subsequent specialization of Mα* in angiosperms enabled the formation of heteromeric Type I MADS TF complexes required for the regulation of endosperm development.

## Conclusion

Angiosperms are the most abundant and diverse group among land plants. The success of angiosperms is closely connected to the developmental innovations of flowers and fruits, as well as the process of double fertilization, coupling fertilization to the formation of the embryo nourishing endosperm tissue (Baroux and Grossniklaus 2019).

Duplication and diversification of type II MADS-box genes underpin the evolution of flowers and fruits in angiosperms (Irish and Litt 2005; Ruelens et al. 2017), while the role of type I MADS-box genes for angiosperm evolution remained obscure. Based on our data, we propose that the origin of the embryo nourishing endosperm tissue is linked to the angiosperm-specific duplication of Type I MADS-box genes (Fig. 4). In the early diverging land plants, ancestral Mα and Mβ/γ-like TFs likely formed heterodimers that had reproductive function based on the expression of gymnosperm Mα and Mβ/γ TFs in female cones and seeds. After the angiosperm lineage diverged from the gymnosperms, true Mγ TFs arose by gene duplication, experienced neofunctionalization, and drove the concerted divergence of some Mα TFs formed by angiosperm-specific gene duplication events. These novel Mγ-Mα heterodimers adopted a function as master regulators of the endosperm developmental network in flowering plants. This proposed scenario is strongly supported by the specific or preferential expression of Mγ and Mα*genes in the endosperm of all sampled angiosperm species as well as functional data in *A.thaliana* and rice, revealing that Mγ and Mα* TFs are required for endosperm development (Chen et al. 2016; Batista et al. 2019; Paul et al. 2020). In contrast to gymnosperms that only have few Type I MADS-box genes (Gramzow et al. 2014); in angiosperms, their number strongly amplified, correlating with the evolution of the embryo-nourishing endosperm. The link between Mγ TFs and endosperm evolution was furthermore supported by the negligible expression of Mγ genes in perispermic seeds, in which the maternal perisperm instead of the endosperm supports embryo growth (Lu and Magnani 2018). The maternal nourishing function in perispermic seeds correlates with the expression of Mβ genes, consistent with the proposed ancestral role of pre-duplicated Mβ/γ genes in regulating nutrient transfer from the maternal tissues to the embryo.

**Fig. 4.**
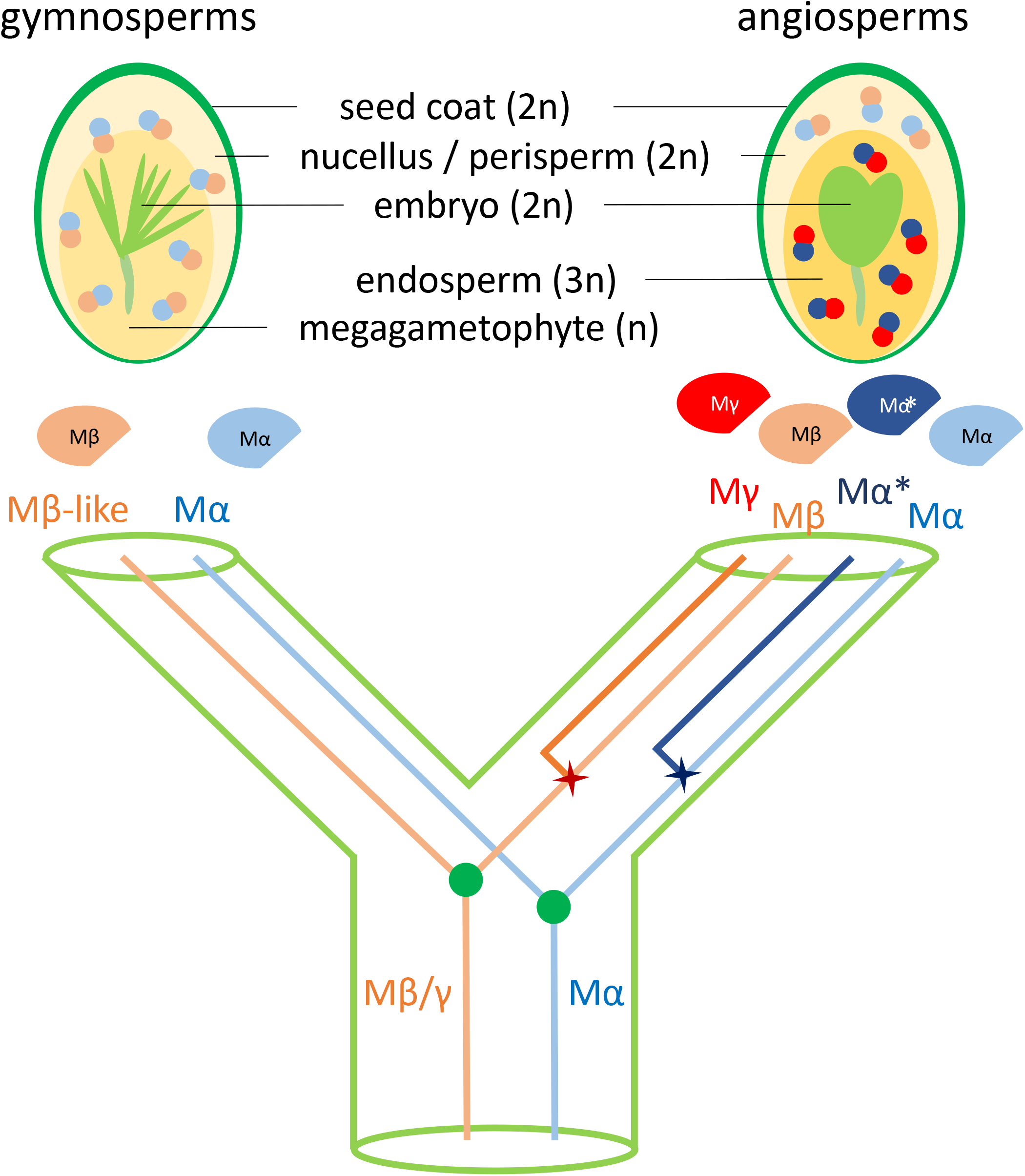
Model depicting the role of Type I MADS-box TFs in the evolution of embryo-nourishing tissues in seed plants. In early seed plants, ancestral Mα and Mβ/γ TFs likely formed heterodimers that regulated the embryo-nourishing behaviour. The gymnosperm Mα and Mβ-like TFs dimerize and function in maternal tissues. Angiosperms express Mα and Mβ heterodimers in maternal tissues, while Mα* and Mγ TFs that arose from angiosperm-specific duplications function in the endosperm and likely enabled endosperm evolution.

Together, our work provides new insights into the role of Type I MADS-box proteins in the origin and evolution of the endosperm, a developmental novelty associated with the rise and diversification of angiosperms.

## Methods

### Phylogenetic analyses

Amino acid sequences of Type I and Type II MADS-box proteins of *A. thaliana* obtained from TAIR10 were used as queries to search for MADS-box proteins in other plant species. The sequences of coding genes in land plant lineages were obtained from PLAZA 4.0 (https://bioinformatics.psb.ugent.be/plaza/; Van Bel et al. 2018), Phytozome v.12 (https://phytozome.jgi.doe.gov/; Goodstein et al., 2012), CoGe (https://genomevolution.org/coge/) or other taxon-themed databases (Supplementary Table S1). MADS-box genes were obtained through reciprocal best BLASTP hits with *A. thaliana* MADS-box genes. The presence of MADS domain in the BLASTP output sequences was further confirmed by the conserved domain search tool, CD-Search (Marchler-Bauer and Bryant 2004) by aligning to the MADS domain entries in the Conserved Domain Database (Lu et al. 2020).

MUSCLE was used to generate the amino acid alignments of MADS-box domains extracted from the identified genes with default settings (Edgar 2004). IQ-TREE 1.6.7 was applied to perform phylogenetic analyses for maximum likelihood trees (Nguyen et al. 2015). The implemented ModelFinder determined LG amino acid replacement matrix (Le and Gascuel 2008) to be the best substitution model in the tree inference (Kalyaanamoorthy et al. 2017). 1000 replicates of ultrafast bootstraps were applied to estimate the support for reconstructed branches (Hoang et al. 2018). The Mα, Mβ and Mγ Type I genes were curated from the phylogenetic position with the defined *Arabidopsis* MADS-box genes.

### Expression analyses

The expression data of Type I MADS-box genes in *A. thaliana* were extracted from Klepicova et al. (2016) for a spectrum of different organ types and developmental stages and Belmonte et al. (2013) for specific compartments in developing seeds. The other transcriptomes used in this study were retrieved from maize (Chen et al. 2014; Walley et al. 2016), rice (Paul et al. 2020), soybean (Chen et al. 2021), castor bean (Xu et al. 2014), tomato (Pattison et al. 2015), coconut (Saensuk et al. 2016), avocado (Ge et al. 2019), monkeyflower (Flores-Vergara et al. 2019), coffee (Ivamoto et al. 2017), *Nymphaea thermarum* (Povilus and Friedman 2021), *Picea* (Nystedt et al. 2013), and *Gnetum* (Hou et al. 2019; Deng et al. 2020).

## Acknowledgements

We thank Dr. Rebecca Povilus and Dr. William Friedman for sharing the seed transcriptome data of *Nymphaea thermarum*. We thank Dr. Qin Li for the comments on the data analyses and visualization. This work was supported by a grant from the Swedish Research Council to CK, a grant from the Knut and Alice Wallenberg Foundation to CK, and support from the Göran Gustafsson Foundation for Research in Natural Sciences and Medicine to CK.

## Author Contributions

Conceptualization, Y.Q. and C.K.; Methodology, Y.Q.; Investigation, Y.Q.; Validation, Y.Q. and C.K.; Formal Analysis, Y.Q.; Data Curation, Y.Q. and C.K.; Writing – Original Draft, Y.Q. and C.K.; Writing – Review & Editing, Y.Q. and C.K.; Visualization, Y.Q.; Funding Acquisition, C.K.; Supervision, C.K.

## Supplementary Figures and Tables

Fig. S1. Phylogeny of Type I MADS-box TFs in several focused groups of land plants with *Arabidopsis thaliana* MADS-box genes (highlighted in red) as references, shown by ML trees with bootstrap values supporting the branches. Gene identifiers in Supplementary Table S1-2. Suffixes of “a”, “b”, “g” denote Mα, Mβ or Mγ TFs, respectively; ‘x’ after gene identifier denotes the second MADS domain in the gene. Genes from selected species were colored to demonstrate the presence of Mγ genes and the two or more clusters of Mα genes. A. ANA, Magnoliids, Ceratophyllales and Proteales; *Amborella trichopoda* (yellow), *Nymphaea thermarum* (purple), *Cinnamomum camphora* (blue). B. Monocots; *Oryza sativa* (yellow), *Zostera marina* (purple), *Asparagus officinalis* (blue). C. Asterids and Caryophyllales; Chrysanthemum nankingense (yellow), Erythranthe guttata (purple), Solanum lycopersicum (blue). D. Rosids; Eucalyptus grandis (yellow), Theobroma cacao (purple), Glycine max (blue).

Fig. S2. Expression of Type I MADS-box genes in reproductive tissues of maize (A), soybean (B) and tomato (C), showing endosperm-expression of Mγ genes. The expression values were normalized into a 0-1 spectrum, with the max value set as 1. A. Gene expression levels at two developmental stages of seed tissues in maize and average expression level in non-endosperm tissues across the whole plant. B. Gene expression levels at two developmental stages of seed tissues in soybean. C. Gene expression levels in two genotypes (LA1589 / *SUN*) of seed and fruit tissues in tomato, before and after fertilization. en/end: endosperm; em/emb: embryo; dap: days after pollination.

Fig. S3. Expression of Type I MADS-box genes in reproductive tissues of *Nymphaea thermarum* (A), *Picea abies* (B) and *Gnetum luofuense* (C), showing Mβ expression. The expression values were normalized into a 0-1 spectrum, with the max value set as 1. A. Gene expression levels at unfertilized ovule and two developmental stages of seeds in *Nymphaea thermarum*; daa: days after anthesis. B. Gene expression levels of female and male cones in *Picea abies* and average expression level in vegetative tissues; veg: vegetative. C. Gene expression levels at mature vs immature seeds in *Gnetum luofuense* and fertile vs sterile ovules.

Table. S1. List of surveyed species.

Table. S2. List of identified Type I MADS-box genes in the surveyed species.

