## Supplemental Figures for "Endosperm evolution by duplicated and neofunctionalized Type I MADS-box transcription factors"

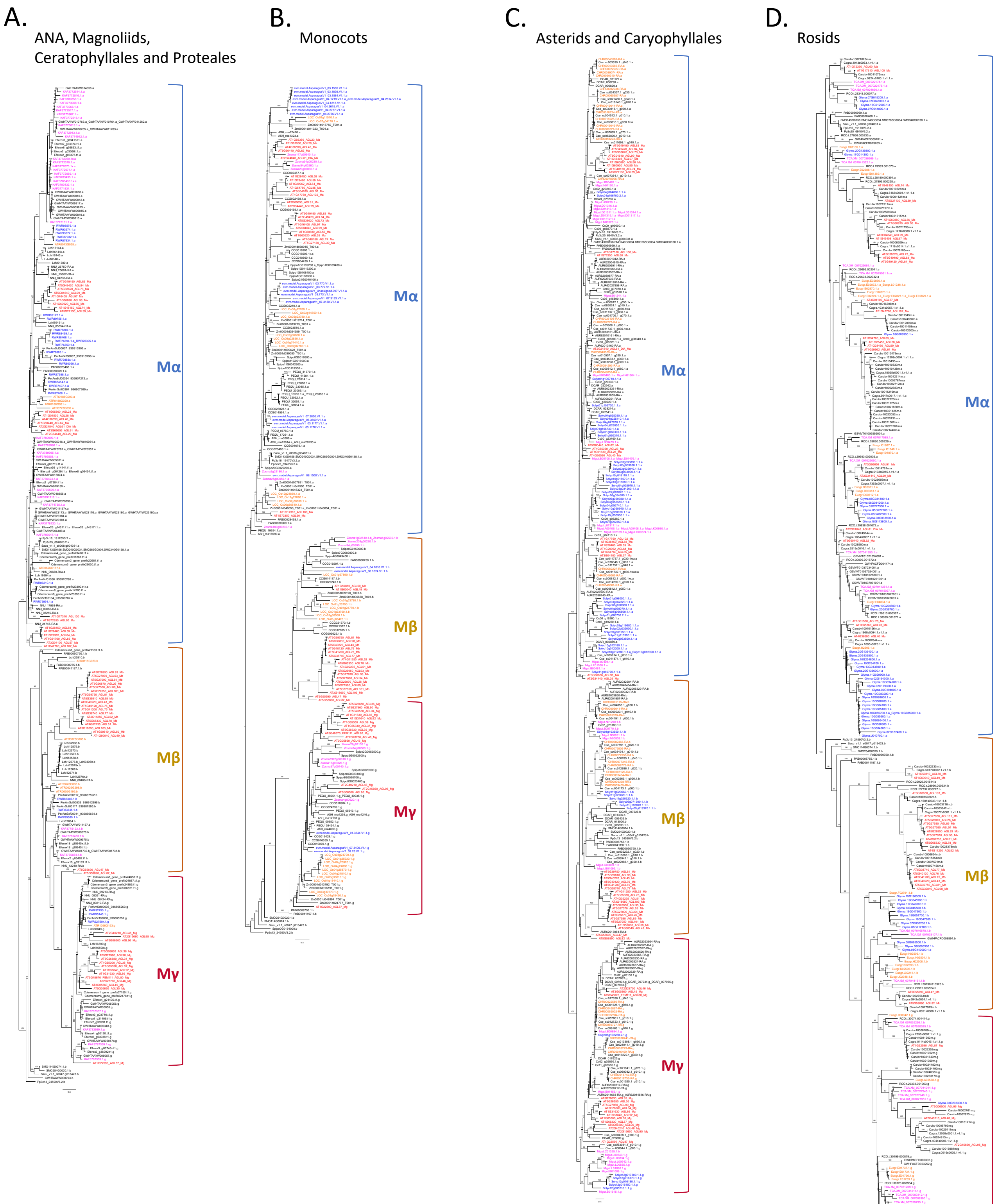


Fig. S1. Phylogeny of Type I MADS-box TFs in several focused groups of land plants with *Arabidopsis thaliana* MADS-box genes (highlighted in red) as references, shown by ML trees with bootstrap values supporting the branches. Gene identifiers in Supplementary Table S1-2. Suffixes of "a", "b", "g" denote Mα, Mβ or Mγ TFs, respectively; 'x' after gene identifier denotes the second MADS domain in the gene. Genes from selected species were colored to demonstrate the presence of Mγ genes and the two or more clusters of Mα genes. A. ANA, Magnoliids, Ceratophyllales and Proteales; *Amborella trichopoda* (yellow), *Nymphaea thermarum* (purple), *Cinnamomum camphora* (blue). B. Monocots; *Oryza sativa* (yellow), *Zostera marina* (purple), *Asparagus officinalis* (blue). C. Asterids and Caryophyllales; *Chrysanthemum nankingense* (yellow), *Erythranthe guttata* (purple), *Solanum lycopersicum* (blue). D. Rosids; *Eucalyptus grandis* (yellow), *Theobroma cacao* (purple), *Glycine max* (blue).

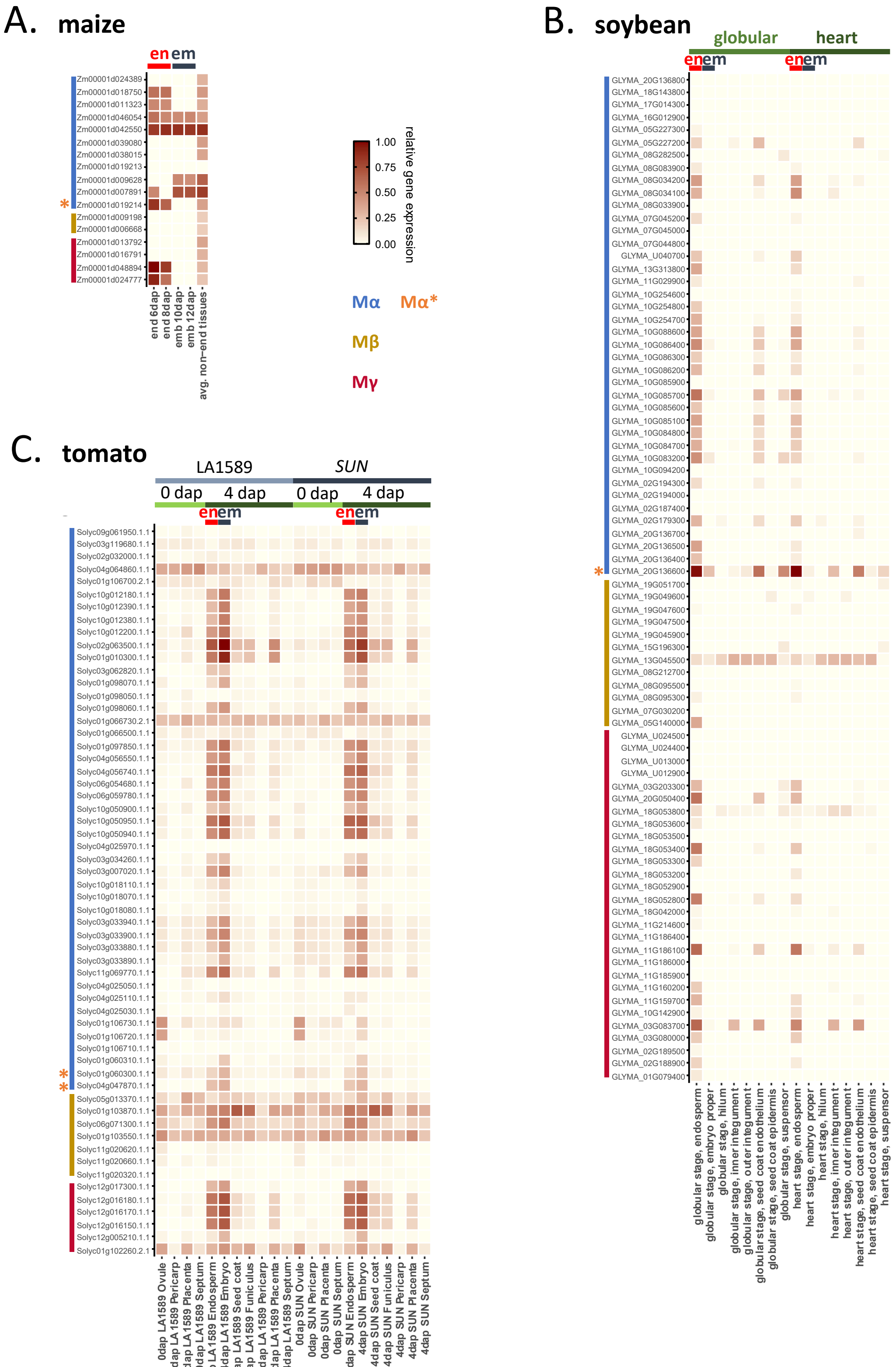


Fig. S2. Expression of Type I MADS-box genes in reproductive tissues of maize (A), soybean (B) and tomato (C), showing endosperm-expression of Mγ genes. The expression values were normalized into a 0-1 spectrum, with the max value set as 1. A. Gene expression levels at two developmental stages of seed tissues in maize and average expression level in non-endosperm tissues across the whole plant. B. Gene expression levels at two developmental stages of seed tissues in soybean. C. Gene expression levels in two genotypes (LA1589 / *SUN*) of seed and fruit tissues in tomato, before and after fertilization. en/end: endosperm; em/emb: embryo; dap: days after pollination.

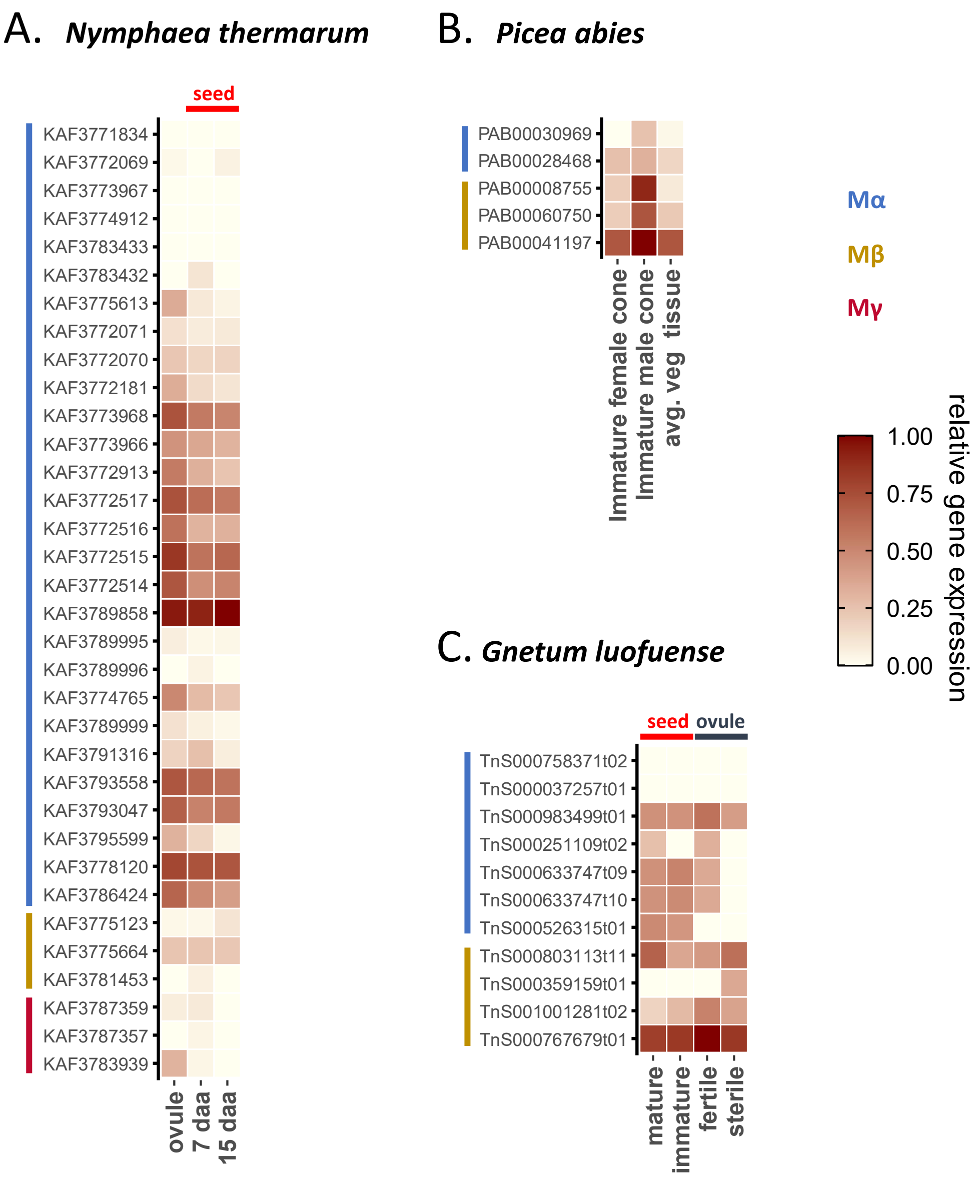


Fig. S3. Expression of Type I MADS-box genes in reproductive tissues of *Nymphaea thermarum* (A), *Picea abies* (B) and *Gnetum luofuense* (C), showing Mβ expression. The expression values were normalized into a 0-1 spectrum, with the max value set as 1. A. Gene expression levels at unfertilized ovule and two developmental stages of seeds in *Nymphaea thermarum*; daa: days after anthesis. B. Gene expression levels of female and male cones in *Picea abies* and average expression level in vegetative tissues; veg: vegetative. C. Gene expression levels at mature vs immature seeds in *Gnetum luofuense* and fertile vs sterile ovules.
